# Propagule and Juvenile-derived Foraminiferal eDNA across intertidal habitats and its implications for accurate sea-level reconstruction

**DOI:** 10.64898/2026.02.24.707652

**Authors:** Zhaojia Liu, Nicole S. Khan, Magali Schweizer, Celia Schunter

**Affiliations:** Department of Earth and Planetary Sciences and the Swire Institute of Marine Science, The University of Hong Kong, Hong Kong SAR; Université d’Angers, Nantes Université, Le Mans Université, CNRS, Laboratoire de Planétologie et Géosciences, LPG UMR 6112, 49000 Angers, France; Swire Institute of Marine Science, School of Biological Sciences, The University of Hong Kong, Hong Kong SAR; State Key Laboratory of Marine Environmental Health and Department of Chemistry, City University of Hong Kong, Tat Chee Avenue, Kowloon, Hong Kong SAR, China

**Keywords:** Environmental DNA, Intertidal, Foraminifera, Propagule, Relative sea level

## Abstract

Foraminiferal environmental DNA (eDNA) assemblages have recently emerged as a robust and complementary proxy for relative sea level (RSL) reconstruction. However, unlike traditional morphological methods, eDNA assemblages are influenced by diverse DNA sources, including propagules and juveniles, whose effects on RSL reconstruction remain poorly understood. To assess how foraminiferal eDNA from different life stages vary in taxa composition and impact RSL reconstruction, we analyzed foraminiferal eDNA from bulk, 500–63 μm and <63 μm size fraction sediments from mangrove and mudflat environments in subtropical Hong Kong. The eDNA assemblages in size-fractioned sediments displayed distinct patterns from those in bulk sediment eDNA across different environments. The propagule and juvenile-derived eDNA <63 μm fraction exhibited a similar community structure to bulk eDNA in mudflat environments but diverged in mangrove environments, indicating a greater contribution of propagule and juvenile eDNA to the total eDNA pool in the mudflat environment. We applied Bayesian transfer function modeling to estimate the elevation of samples using different size fractions. eDNA assemblages from the <63 μm fraction systematically underpredicted elevation in mangrove environments, while elevations inferred from the 500–63 μm fraction and bulk sediment eDNA were accurate. Conversely, all eDNA assemblages in the mudflat-mangrove transitional zone led to the overprediction of RSL. These findings confirm the reliability of bulk sediment eDNA for RSL reconstruction in mangrove environments, while highlighting the need for caution when reconstructing RSL in transitional zones.

## 1. Introduction

High-resolution reconstructions of relative sea level (RSL) are essential for understanding coastal evolution and the drivers of RSL change (Horton et al., 2006; Kemp et al., 2013; Khan et al., 2019; Patterson et al., 2005; Williams et al., 2023; Yu et al., 2025). Among geological proxies, foraminifera are one of the most widely used sea-level indicators due to their well-defined and quantifiable relationship with tidal elevation (Edwards & Horton, 2000; Rush et al., 2021; Williams et al., 2023), allowing for high-resolution (sub-decimeter) reconstruction of RSL changes where foraminifera are preserved within continuous salt-marsh archives (Barnett et al., 2016; Rossi et al., 2011; Rush et al., 2021). Traditional morphology-based RSL reconstructions typically construct modern training sets from dead adult assemblages in the 63–500 µm size range to facilitate identification and minimize temporal variability (Hayward et al., 1996; Horton & Edwards, 2006; Kemp et al., 2011; Murray & Alve, 2000; Scott et al., 2007; Walker et al., 2020).

While morphology-based methods have demonstrated considerable success in reconstructing past sea levels, they remain constrained by incomplete preservation of foraminiferal tests and challenges in distinguishing morphologically similar taxa (Berkeley et al., 2007; Khan et al., 2019; Yu et al., 2025). Environmental DNA (eDNA) has emerged as a powerful tool in foraminiferal ecological research (Brinkmann et al., 2023; He et al., 2019; Pawlowski et al., 2016; Singer et al., 2023) that circumvents these constraints by not relying on the identification of tests. Recent work has demonstrated that bulk sediment-derived foraminiferal eDNA assemblages exhibit clear vertical zonation across tidal elevation gradients, extending the utility of eDNA as a complementary proxy for late Holocene RSL reconstruction (Liu et al., 2025a). Compared to traditional morphological methods, the eDNA approach detects both fossilizing and non-fossilizing taxa, offering a comprehensive picture of community composition (Lejzerowicz et al., 2013; Pawłowska et al., 2014; Singer et al., 2023), which may improve the resolution of RSL reconstructions (Liu et al., 2025a). The preserved eDNA in the sediment was detected even where shells were no longer preserved, giving it additional utility as a complementary tool for paleoenvironmental reconstruction (Liu et al., 2025a).

However, this sensitivity of eDNA to various DNA sources also presents challenges when interpreting the relationship between foraminiferal eDNA assemblages and tidal elevation (Liu et al., 2025a). As single-celled protists, benthic foraminifera reproduce both sexually and asexually, producing highly mobile propagules and juveniles that disperse rapidly across intertidal environments (Alve & Goldstein, 2002, 2003; Goldstein & Alve, 2011). In contrast to morphological assemblages, foraminiferal eDNA assemblages not only contain the local adult population, but also inevitably incorporate extracellular DNA sources (autochthonous or allochthonous) and DNA from propagules and juveniles (Ellegaard et al., 2020; Taberlet et al., 2012). This propagule and juvenile-derived eDNA can contribute significantly to modern coastal sediments (Brinkmann et al., 2023; Singer et al., 2023). The high mobility of foraminiferal propagules and juveniles may broaden the apparent elevation range of eDNA assemblages, potentially introducing uncertainty and bias into RSL reconstructions. The extent to which propagule and juvenile DNA contribute to foraminiferal eDNA assemblages, and its influence on the development of sea-level proxies, remains poorly understood.

To address this knowledge gap, we analyzed foraminiferal eDNA assemblages in different size-fractions of sediments across subtropical mudflat and mangrove environments in Hong Kong. Specifically, this study aims to: (1) investigate compositional differences in foraminiferal eDNA assemblages among 500-63 μm, <63 μm, and bulk sediment fractions across different environments, and (2) assess how <63 μm fraction eDNA (propagule/juvenile-dominated) influences the accuracy of eDNA-based RSL reconstructions. Our results demonstrate the influence of foraminiferal propagules and juveniles on eDNA assemblages in coastal wetlands and highlight important considerations for their application in paleoenvironmental and RSL reconstruction.

## 2. Methods

### 2.1 Study area

Mai Po Nature Reserve, situated in southeastern Deep Bay within the Pearl River Estuary (Fig. 1B), was chosen as the study area. This subtropical area is characterized by hot, humid summers (maximum average monthly temperature ∼30°C in July) and mild winters (minimum average monthly temperature ∼17°C in January). Annual rainfall totals 2,775 mm (2023), 77% of which occurs during the wet season (April-September) (HongKong-Observatory, 2024). Deep Bay’s brackish conditions result from freshwater input from the Pearl and Shenzhen Rivers. The bay is relatively shallow, with an average water depth of approximately 2.9 m (Li & Lee, 1998) and semidiurnal tides, with an amplitude of 1.96 m from mean higher high water (MHHW) to mean lower low water (MLLW), as measured at the Tsim Bei Tsui gauge (Fig. 1C).

**Fig. 1.**
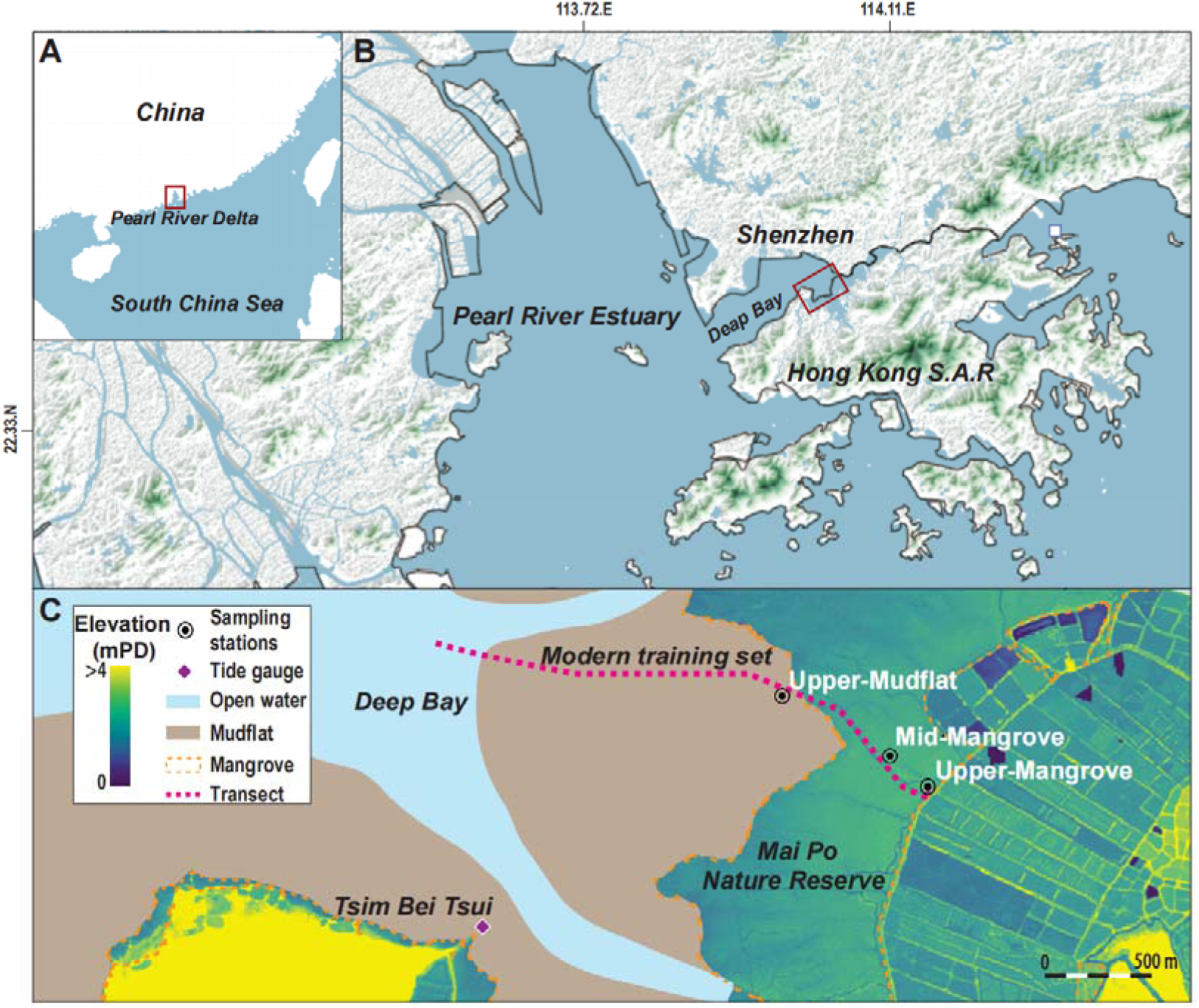
Location of the study area in the Pearl River Delta (A) and Deep Bay (B). (C) Location of upper-mudflat, mid-mangrove, and upper-mangrove sampling stations within the Mai Po Nature Reserve. The position of surface transects of established modern training set in Liu et al (2025a) are also shown. The color map shows land surface elevation derived from the LiDAR digital elevation model provided by the Hong Kong S.A.R. government, with the location of nearest tide gauge at Tsim Bei Tsui also shown.

Sampling was conducted in November 2023, during the dry season (with average temperature of 23.5°C and monthly total precipitation of 3.3 mm)—a period when monothalamids are reproducing and dominate the foraminiferal eDNA assemblages (Liu et al., 2025b). Three sampling stations were selected to capture compositional differences in eDNA assemblages across an intertidal gradient: (1) mudflat-mangrove transitional zone (upper-mudflat station), (2) mangrove forest (mid-mangrove station), and (3) interior mangrove forest (upper-mangrove station) of Mai Po Nature Reserve. These stations coincided with established seasonal monitoring stations (Fig. 1C) (Liu et al., 2025b), which were originally selected to capture the seasonality of eDNA assemblages along the seaward-landward environmental gradient during one and half year. Managed by the World Wide Fund for Nature (WWF) since 1975, Mai Po Nature Reserve comprises extensive intertidal mudflats and mangrove ecosystems. The dominant mangrove species are *Kandelia candel*, *Aegiceras corniculatum*, and *Avicennia marina* (Lee, 2000; Li et al., 2019). The vegetation in the mudflat–mangrove transitional zone was dominated by *Sporobolus alterniflorus*, which progressively encroached upon and eventually covered the originally barren monitoring station during the study period (Liu et al., 2025b), with occasional occurrences of the non-native species *Sonneratia apetala* (Yu et al., 2025). The three sampling stations were situated adjacent to a foraminiferal eDNA modern training set previously established along a transect within the Mai Po Nature Reserve (Fig. 1C) (Liu et al., 2025b).

### 2.2 Sampling strategy

At each sampling station, three replicate samples were collected randomly from an approximately 1 × 1 m² plot for eDNA analysis to maximize the recovery of genetic diversity (Brinkmann et al., 2023; Singer et al., 2023). To provide context for comparing eDNA assemblages in different size fraction, at each station, three replicate samples for morphological analysis were also collected directly adjacent to the eDNA replicates. The sampling procedure followed protocols detailed in Liu et al (2025a; 2025b). Briefly, approximately 20 cm^3^ from the top 1 cm of surface sediment was sampled for eDNA and morphological analyses. eDNA collection was performed using sterile, disposable or 10% bleach-sanitized equipment and transferred into a 50 ml Falcon tube to prevent cross-contamination. eDNA samples were kept on ice during fieldwork and stored at –20°C immediately upon return to the laboratory on the sampling day to minimize DNA degradation Samples for morphological analysis were placed in 50 ml Falcon tubes, and preserved in an 50% ethanol-calcite buffer solution. Samples were stained immediately with rose Bengal after collection in the field to distinguish living from dead foraminifera (Walton, 1952). These samples were transported in a cool box on ice and stored in 4°C refrigerator in the laboratory to preserve specimen integrity.

The elevation of each sampling station was first measured relative to a temporary local benchmark using a total station, and then referenced to the Hong Kong Principal Datum (PD) with a Leica GS18 GNSS system. Elevation was measured three times due to the slightly uneven surface of the sampling stations. These elevations were subsequently converted to the standardized water level index units (SWLI) (Kemp & Telford, 2015), which assigns a value of 100 to mean tide level (MTL) and 200 to mean higher high water (MHHW), using tidal datums recorded by the nearby Tsim Bei Tsui tide gauge (Fig. 1C).

### 2.3 Sample processing and eDNA extraction

To separate eDNA from different sediment size fractions, 9 collected samples were wet-sieved using autoclaved Milli-Q water through 500 and 63 μm mesh sieves. Additionally, to assess whether using samples from barren mudflat rather than from the transitional-zone mudflat affects sea-level estimation, six additional samples (three from mangrove, one from the transitional zone, and two from barren mudflat) collected in Liu et al. (2025a) were also processed with wet-sieving. The <63 μm fraction was expected to be enriched with potential small unkown species, juveniles and propagules (Alve & Goldstein, 2014), while the 500–63 μm fraction was expected to represent the conventional adult foraminiferal DNA pool. All equipment (e.g., sieves) was thoroughly sterilized prior to use by: (1) sonication for 15 minutes, (2) rinsing with soapy water, (3) immersion in 10% bleach for ∼60 seconds, and (4) a final Milli-Q water rinse to eliminate potential contaminants. After sieving, obtained liquid containing the <63 μm fraction was collected into 50 mL Falcon tubes. The <63 μm fraction samples were subsequently filtered through sterile 0.47 μm glass fiber filters to capture and enrich the eDNA (Bessey et al., 2022). Both the filters (containing the <63 μm fraction) and the retained 500–63 μm sediment fractions were stored in 50 mL Falcon tubes at –20 °C until further processing. eDNA was extracted from 1) the glass fiber filters (<63 μm fraction), 2) subsamples of the 500–63 μm sediment fraction, and 3) bulk sediment, using the 0.5g-PowerSoil® DNA Isolation Kit (QIAGEN) following manufacturer protocol in a sterile laboratory environment. Extracted DNA samples were then stored at –80°C prior to downstream molecular analyses.

### 2.4 PCR metabarcoding and sequencing

We performed a two-step PCR amplification following methodologies from Liu et al (2025a; 2025b). Briefly, each DNA extraction was amplified in technical duplicate. The first PCR amplified the 135-190 bp 18S rDNA fragment using primers s14F1-s15 with Illumina adapters (Pawlowski et al., 2014) following Illumina MiSeq System sequencing protocol in 25 µL reactions containing 12.5 μl of Taq PCR Master Mix (QIAGEN), 4 µl of each primers (1 μM), 1.25 µl Mg² □ (1 μM), 0.75 μl Mill-Q water and 2.5 μl template DNA. PCR cycling conditions were as follows: an initial denaturation at 94 °C for 3 minutes; 45 cycles consisting of 94 °C for 30 seconds, 55 °C for 30 seconds, and 72 °C for 45 seconds; and a final extension at 72 °C for 5 minutes. Replicate PCR products from each sample were pooled and purified using AMPpure XP beads (Beckman Coulter, Singapore) following the Illumina MiSeq System sequencing protocol. Next, an indexing PCR was performed with the Nextera XT Index Kit (Illumina) to attach dual indices to the purified amplicons. Each indexing PCR reaction (50 µL total volume) contained 25 µL of 2× KAPA HiFi HotStart ReadyMix, 5 µL of each index primer (10 µM), 10 µL of PCR-grade water, and 5 µL of purified PCR product. The cycling conditions for the indexing PCR included an initial denaturation at 95 °C for 3 minutes; 8 cycles of 95 °C for 30 seconds, 55 °C for 30 seconds, and 72 °C for 30 seconds; followed by a final extension at 72 °C for 5 minutes. Indexed PCR products were purified again using AMPpure XP beads. Finally, purified samples were pooled to a final concentration of 20 μM in 10 mM Tris buffer (pH 8.5) and submitted to Novogene Co., Ltd for sequencing on the NovaSeq System platform with paired-end 150 bp reads (PE 150).

### 2.5 Bioinformatic processing

Bioinformatics processing followed the pipeline described in Liu et al (2025a; 2025b). In brief, raw sequence data were demultiplexed and primers removed using Cutadapt v2.8 (Martin, 2011). The resulting paired-end reads were merged and quality-trimmed with PEAR v0.9.11, applying a quality threshold of 26 (Zhang et al., 2014). Concatenated sequences were then denoised and chimeras removed using VSEARCH (Rognes et al., 2016). Sequences were clustered into operational taxonomic units (OTUs) at 97% similarity with VSEARCH. Quality control filters excluded OTUs shorter than 90 bp, those with fewer than 10 reads, or those detected in negative controls with read counts matching or exceeding their counts in any sample. Taxonomic assignment was performed using BLASTn v2.12.0 (Altschul et al., 1990) referring to the GenBank nucleotide database and an in-house foraminiferal DNA database, with an e-value cutoff of 0.001. Assignments were made using python script taxonomy_assignment_BLAST_V1.py (Joseph7e, 2020) at species, family, and phylum levels based on identity thresholds of 98%, 90%, and 80%, respectively. OTUs with identities between 80% and 90% (assigned at the phylum level) and those with unknown taxonomy were classified as “undetermined OTUs” and treated as individual taxon in subsequent analyses. OTUs with identity below 80% were excluded as non-foraminiferal sequences.

### 2.6 Morphological foraminiferal analysis

To provide context for the eDNA foraminiferal assemblages, we also conducted morphological analyses at each sampling station following procedure detailed in Liu et al (2025a). Briefly, foraminiferal tests were isolated by wet-sieving sediment samples to isolate the 63–500 µm fraction. Sieved samples were wet-split to divide samples into eight equal parts (Horton, 1997; Horton & Edwards, 2006; Scott & Hermelin, 1993), followed by manual picking under a stereoscope (Callard et al., 2011; Hawkes et al., 2010; Lal et al., 2020). Taxonomic identification was based on published references from our study area (Liu et al., 2025a; Yu et al., 2025). Where species-level identification was uncertain due to poor preservation of diagnostic test features, specimens from the genera *Ammobaculites* and *Miliammina*, as well as the species *Ammonia tepida*, *Ammonia convex, Ammonia beccarii* and *Ammonia confertitesta*, were grouped as *Ammobaculites* spp., *Miliammina* spp., and *Ammonia* spp., respectively (Culver & Horton, 2005; Kemp et al., 2009; Khan et al., 2019; Saunders, 1958; Yu et al., 2025). Living individuals were determined by the presence of rose Bengal staining on more than one chamber (Horton, 1999; Horton & Edwards, 2003).

### 2.7 Statistical analysis

To assess differences in the abundance of dominant taxa among sediment size fractions and bulk samples at each sampling station, we conducted a one-way analysis of variance (ANOVA; 1000 permutations, p value threshold = 0.05) (Clark et al., 1992; Troth et al., 2021). Dominant taxa were defined as those exhibiting >5% relative abundance in at least one sample (Liu et al., 2025a). Before performing ANOVA, we tested for homogeneity of variances for each taxon’s abundance at each station and applied log transformation where necessary to satisfy ANOVA assumptions.

Taxonomic diversity across size fractions and stations were visualized by generating a heatmap of dominant taxa. Replicate samples from each examined size fraction and bulk samples at each station were pooled to provide a representative overview of the taxonomic composition and diversity of eDNA assemblages across different size fractions and stations. The heatmap was generated using the ggheatmap function in the heatmaply R package (Galili et al., 2018).

To quantitatively explore differences in assemblage composition between stations and size fractions, we used non-metric multidimensional scaling (NMDS) based on Bray-Curtis dissimilarity matrices of relative abundance data, with replicate samples included individually, implemented using the vegan package (v2.6-4) in R (Oksanen et al., 2007). The resulting ordination plot displays the first two NMDS dimensions, illustrating compositional variation among stations. To determine whether differences in sediment size fraction significantly contributed to sample clustering, we performed a permutational multivariate analysis of variance (PERMANOVA) with 999 permutations, also based on Bray-Curtis distances (Anderson, 2014). To minimize the disproportionate influence of Family Saccamminidae on assemblage pattern, an additional NMDS was performed on the eDNA dataset after excluding this taxon, enabling clearer visualization of signals from other taxa.

To evaluate how eDNA assemblages from different sediment size fractions at different environments influence RSL reconstructions, we applied a Bayesian transfer function (BTF) (Cahill et al., 2016) to estimate elevations for 63–500 µm, <63 µm size fraction and bulk eDNA sample from 9 samples collected at 3 sampling stations. In addition, we applied the BTF to the 63–500 µm and <63 µm size fractions of six supplementary samples collected in Liu et al. (2025a). The BTF was constructed using the modern training developed in Liu et al (2025a). To account for discrepancies in sequencing depth among samples, eDNA sequence counts were rarefied to match the lowest sequencing depth in the dataset (Hughes & Hellmann, 2005), following the exclusion of one modern sample with a low number of reads. Only taxa with >5% relative abundance in at least one sample were included to reduce bias introduced by rare taxa (Patterson & Fishbein, 1989).

## 3. Results

### 3.1 eDNA assemblage

Saccamminidae dominated eDNA assemblages across all size fractions and stations (Fig. 2A), with relative abundances of 59-80% in the <63 µm fraction across all stations (Fig. S1). Statistical differences among size fractions varied by station: no significant differences were detected at upper-mangrove and mid-mangrove stations (Table 1), while the upper-mudflat station showed significant variation among fractions (ANOVA; *p* <0.05; Table 1), with higher abundances in bulk (45-60%) and <63 µm (59-80%) fractions compared to the 500-63 µm fraction (22-47%).

**Fig. 2.**
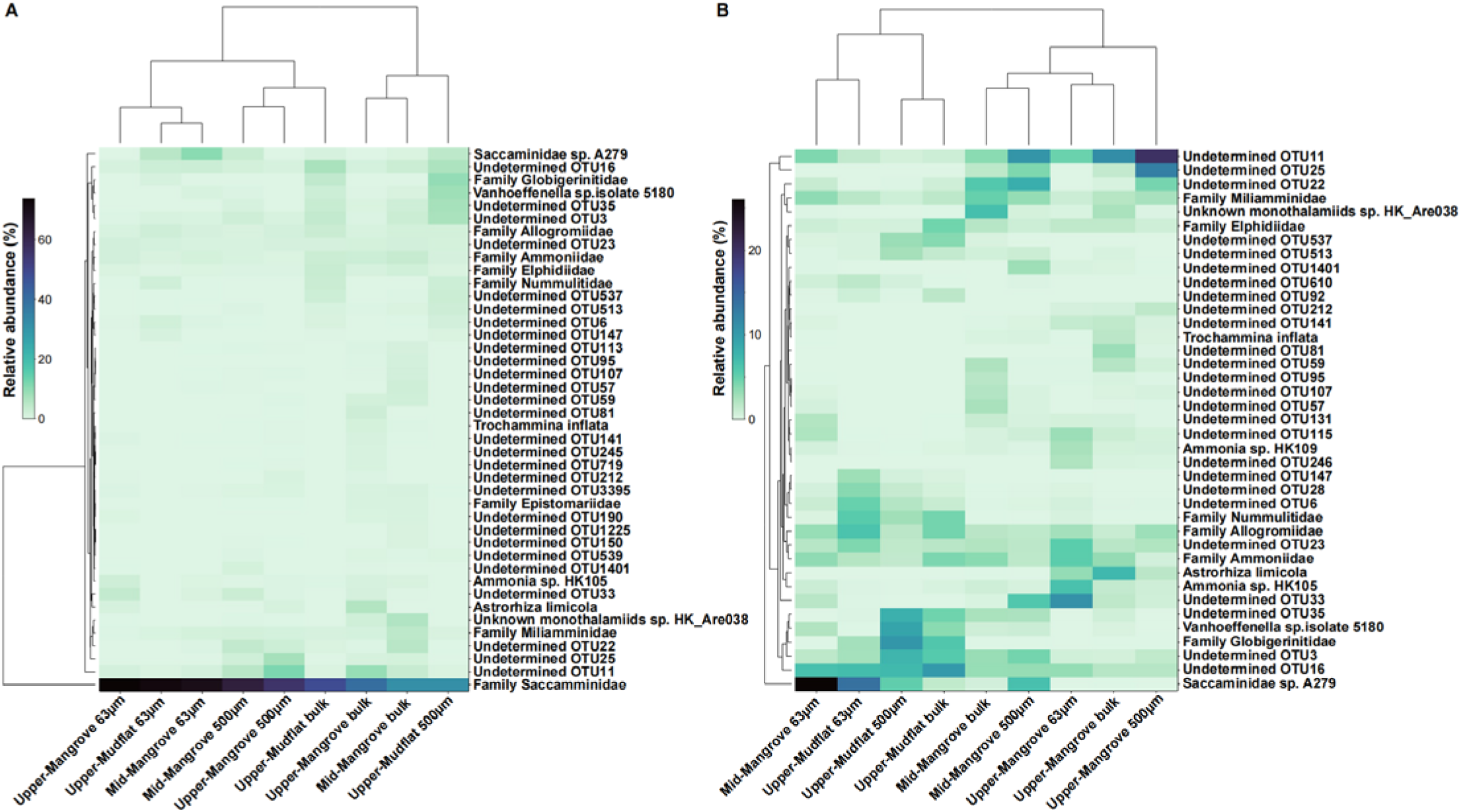
Heatmap showing the relative abundance of dominant foraminiferal taxa at each sampling station across different size fractions and bulk eDNA sample (A) with and (B) without inclusion of Family Saccaminidae, derived from eDNA metabarcoding using foraminifera-specific primers.

**Table 1.**
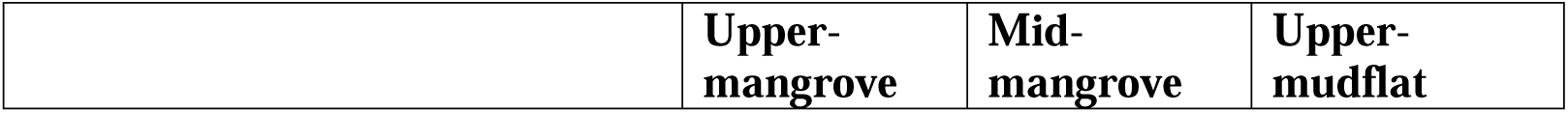
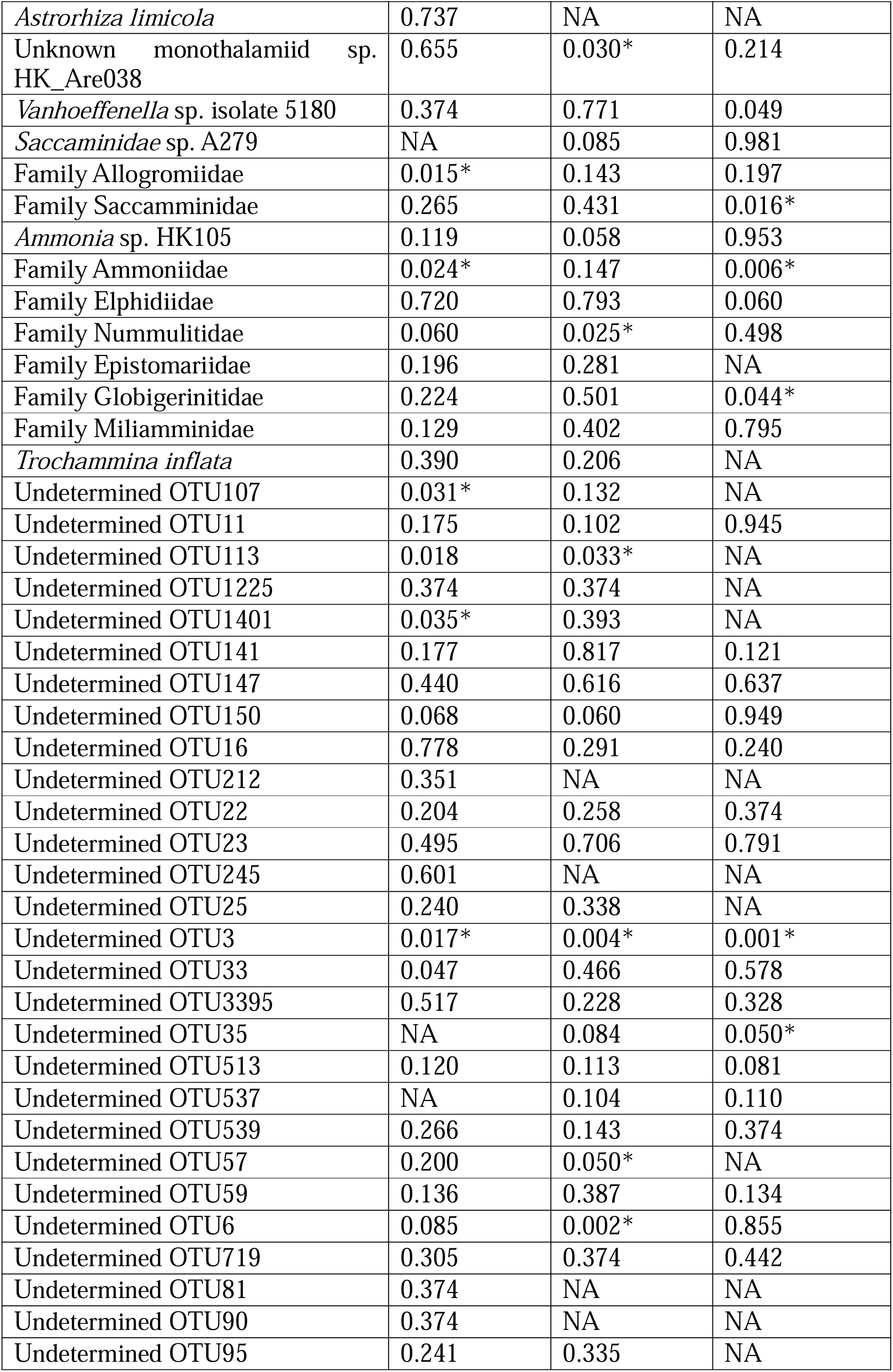
One-way analysis of variance (ANOVA) results of examined dominant taxa (relative abundance >5% in at least one sample). P-value > 0.05 is significant and marked with *. NA indicates that the corresponding taxon at a given station could not be included in the ANOVA because it was absent at that station.

|  | Upper-mangrove | Mid-mangrove | Upper-mudflat |
| --- | --- | --- | --- |
| *Astrorhiza limicola* | 0.737 | NA | NA |
| Unknown monothalamiid sp.<br>HK_Are038 | 0.655 | 0.030* | 0.214 |
| *Vanhoeffenella* sp. isolate 5180 | 0.374 | 0.771 | 0.049 |
| *Saccaminidae* sp. A279 | NA | 0.085 | 0.981 |
| Family Allogromiidae | 0.015* | 0.143 | 0.197 |
| Family Saccaminidae | 0.265 | 0.431 | 0.016* |
| *Ammonia* sp. HK105 | 0.119 | 0.058 | 0.953 |
| Family Ammoniidae | 0.024* | 0.147 | 0.006* |
| Family Elphidiidae | 0.720 | 0.793 | 0.060 |
| Family Nummulitidae | 0.060 | 0.025* | 0.498 |
| Family Epistomariidae | 0.196 | 0.281 | NA |
| Family Globigerinitidae | 0.224 | 0.501 | 0.044* |
| Family Miliamminidae | 0.129 | 0.402 | 0.795 |
| *Trochammina inflata* | 0.390 | 0.206 | NA |
| Undetermined OTU107 | 0.031* | 0.132 | NA |
| Undetermined OTU11 | 0.175 | 0.102 | 0.945 |
| Undetermined OTU113 | 0.018 | 0.033* | NA |
| Undetermined OTU1225 | 0.374 | 0.374 | NA |
| Undetermined OTU1401 | 0.035* | 0.393 | NA |
| Undetermined OTU141 | 0.177 | 0.817 | 0.121 |
| Undetermined OTU147 | 0.440 | 0.616 | 0.637 |
| Undetermined OTU150 | 0.068 | 0.060 | 0.949 |
| Undetermined OTU16 | 0.778 | 0.291 | 0.240 |
| Undetermined OTU212 | 0.351 | NA | NA |
| Undetermined OTU22 | 0.204 | 0.258 | 0.374 |
| Undetermined OTU23 | 0.495 | 0.706 | 0.791 |
| Undetermined OTU245 | 0.601 | NA | NA |
| Undetermined OTU25 | 0.240 | 0.338 | NA |
| Undetermined OTU3 | 0.017* | 0.004* | 0.001* |
| Undetermined OTU33 | 0.047 | 0.466 | 0.578 |
| Undetermined OTU3395 | 0.517 | 0.228 | 0.328 |
| Undetermined OTU35 | NA | 0.084 | 0.050* |
| Undetermined OTU513 | 0.120 | 0.113 | 0.081 |
| Undetermined OTU537 | NA | 0.104 | 0.110 |
| Undetermined OTU539 | 0.266 | 0.143 | 0.374 |
| Undetermined OTU57 | 0.200 | 0.050* | NA |
| Undetermined OTU59 | 0.136 | 0.387 | 0.134 |
| Undetermined OTU6 | 0.085 | 0.002* | 0.855 |
| Undetermined OTU719 | 0.305 | 0.374 | 0.442 |
| Undetermined OTU81 | 0.374 | NA | NA |
| Undetermined OTU90 | 0.374 | NA | NA |
| Undetermined OTU95 | 0.241 | 0.335 | NA |

Foraminiferal eDNA assemblages showed station-specific taxonomic patterns that varied by size fraction. Mangrove stations (upper- and mid-) were characterized by hard-shelled taxa (Miliamminidae 1–7%, Ammoniidae 1–6%) and specific OTUs in coarse fractions (OTU11 ≤23%, OTU25 ≤ 22%), while the upper-mudflat was distinguished by soft-walled monothalamids (Vanhoeffenella sp. 6 – 15%) and morphologically undetectable Globigerinitidae (6–16%) (Fig. 2).

NMDS ordination revealed that size-fraction effects were station-dependent and masked by dominant Saccamminidae. Fine fractions (<63 µm) clustered tightly across all stations due to consistent Saccamminidae dominance, while coarse fractions (500–63 µm) and bulk samples were more dispersed (Fig. 3A). Removing Saccamminidae revealed clearer patterns: upper-mangrove fractions converged on the ordination space while other stations remained distinct (Fig. 3B). PERMANOVA confirmed significant station differences (F = 8.75, R² = 0.42, *p* <0.001) and contrasting size-fraction effects—significant in mid-mangrove (F = 3.18, *p* = 0.029) and upper-mudflat (F = 6.86, *p* = 0.003) but not upper-mangrove (F = 2.04, *p* = 0.12).

**Fig. 3.**
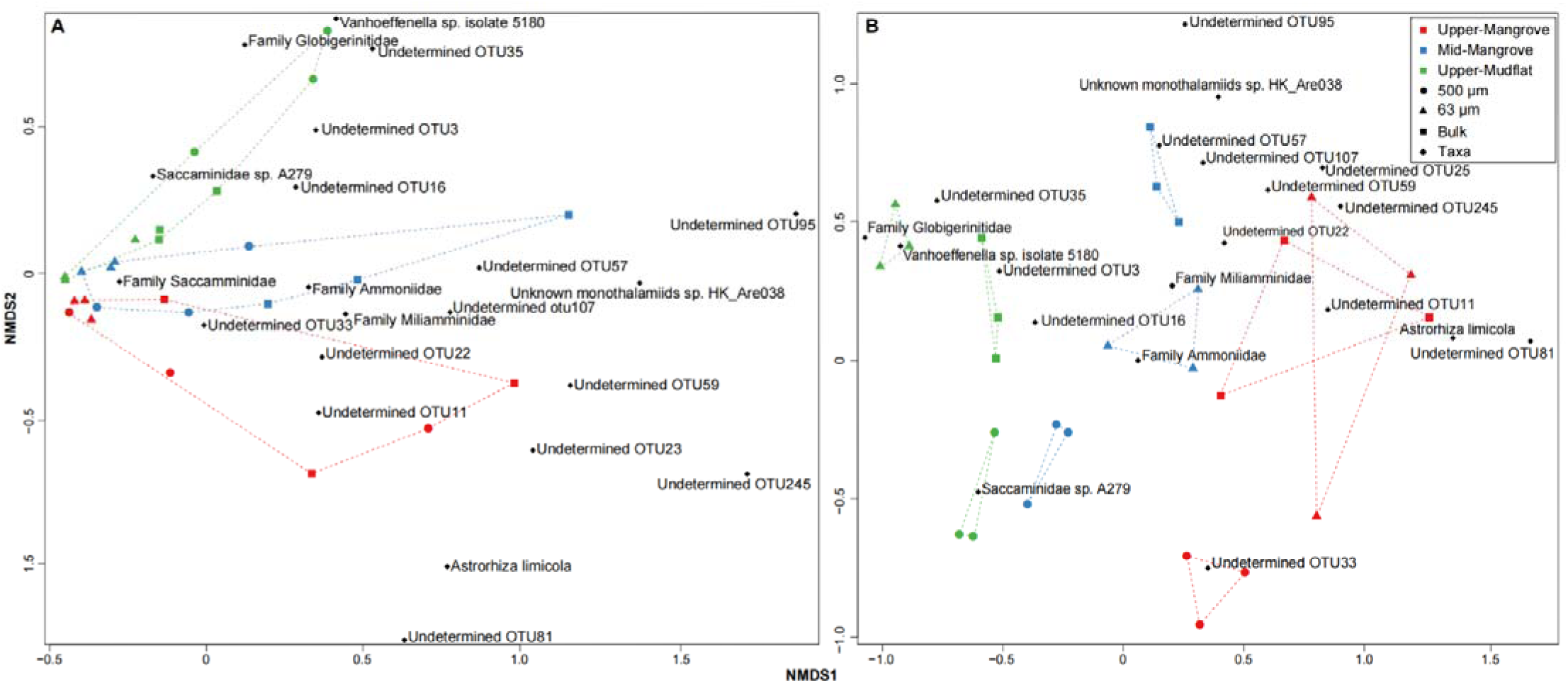
Non-metric Multi-Dimensional Scaling (NMDS) plots based on Bray-Curtis dissimilarities of foraminiferal eDNA assemblages from 500–63 □µm, <63 □µm size fractions, and bulk samples at each sampling station: (A) with and (B) without inclusion of Family Saccaminidae. The NMDS is constructed with the dominant taxa that with >5% relative abundance in at least one sample. Samples collected from different environments and with different size fractions are marked in different colors and shapes.

### 3.2 Morphological assemblage

Morphology-based assemblages displayed patterns distinct from the eDNA data, with station-specific taxonomic compositions dominated by calcareous taxa (Table 2; Fig. S2; Table S1;). Upper-mangrove stations were characterized by moderate Ammoniidae dominance (53–65%) with significant agglutinated contributions (Trochamminidae 4–38%, Miliamminidae 6–22%), while the mid-mangrove showed dominance of Ammoniidae (49–53%) and Miliamminidae (40–44%). The upper-mudflat was distinguished by strong Ammoniidae dominance (68–80%) and reduced Miliamminidae presence (9–26%). The number of stained individuals was consistently low in all stations, though slightly elevated abundance of living Ammoniidae were observed at the upper-mangrove station (Table 3; Table S2).

**Table 2.**
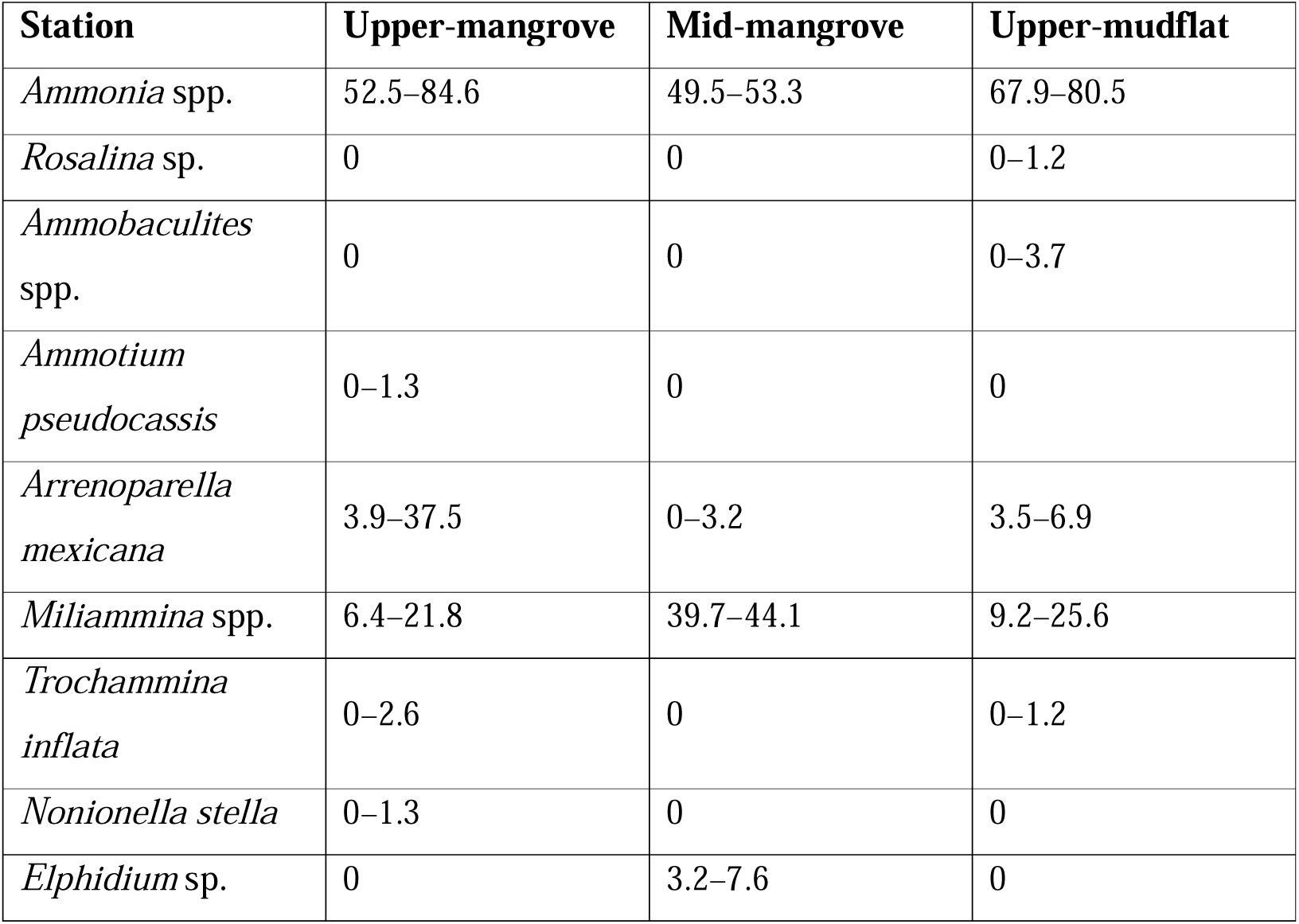
Relative abundance (%) of species for foraminiferal morphological assemblages at each sampling stations.

| Station | Upper-mangrove | Mid-mangrove | Upper-mudflat |
| --- | --- | --- | --- |
| *Ammonia* spp. | 52.5–84.6 | 49.5–53.3 | 67.9–80.5 |
| *Rosalina* sp. | 0 | 0 | 0–1.2 |
| *Ammobaculites* spp. | 0 | 0 | 0–3.7 |
| *Ammotium pseudocassis* | 0–1.3 | 0 | 0 |
| *Arrenoparella mexicana* | 3.9–37.5 | 0–3.2 | 3.5–6.9 |
| *Miliammina* spp. | 6.4–21.8 | 39.7–44.1 | 9.2–25.6 |
| *Trochammina inflata* | 0–2.6 | 0 | 0–1.2 |
| *Nonionella stella* | 0–1.3 | 0 | 0 |
| *Elphidium* sp. | 0 | 3.2–7.6 | 0 |

**Table 3.** Relative abundance (%) of rose Bengal-stained individuals of each taxon against total counts of dead and alive morphological assemblage at each sampling station.

| **Station** | **Upper-mangrove** | **Mid-mangrove** | **Upper-mudflat** |
| --- | --- | --- | --- |
| *Ammonia* spp. | 12.9–17.53 | 5.6–8.8 | 3.3–11.9 |
| *Rosalina* sp. | 0 | 0 | 0 |
| *Ammobaculites* spp. | 0 | 0 | 0 |
| *Ammotium pseudocassis* | 0 | 0 | 0 |
| *Arrenoparella mexicana* | 0–2.1 | 0–0.7 | 0–1.1 |
| *Miliammina* spp. | 0–1.0 | 0–2.1 | 1.0–3.2 |
| *Trochammina inflata* | 0 | 0 | 0 |
| *Nonionella stella* | 0 | 0 | 0 |
| *Elphidium* sp. | 0 | 0 | 0 |

### 3.3 Relative sea level estimation of different size fractions for eDNA and morphological approaches

BTF elevation estimates showed station-dependent patterns in accuracy across size fractions (Fig. 4). At upper- and mid-mangrove stations (observed elevations: 2.17–2.22 m PD, 189–195 SWLI), all samples fell within observed elevation ranges at 95% confidence, although <63 µm fractions consistently underpredicted elevations. Conversely, at the upper-mudflat station (1.27–1.31 m PD, 93–99 SWLI), all size fractions exhibited systematic overprediction, with only the <63 µm fraction capturing observed elevations within 2σ uncertainty. The precision of elevation predictions also varied among fractions and stations. The 500–63 µm fraction from the upper-mudflat station exhibited the narrowest prediction range (99–145 SWLI) and the lowest average 1σ uncertainty (8.5 SWLI). For the additional sieved samples obtained from Liu et al. (2025a), all samples—including both the 500–63 µm and <63 µm fractions—fell within the observed elevation ranges at 95% confidence, except for the sample (22/MPa/42) collected from the transitional zone between mangrove and mudflat, for which both fractions showed underprediction (Fig. S3). The elevation estimated by morphological assemblages accurately captured the observed elevation within 95% uncertainty interval at all three sampling stations, but exhibit greater uncertainties in the upper-mudflat station (45-53 SWLI) (Fig. 4).

**Fig. 4.**
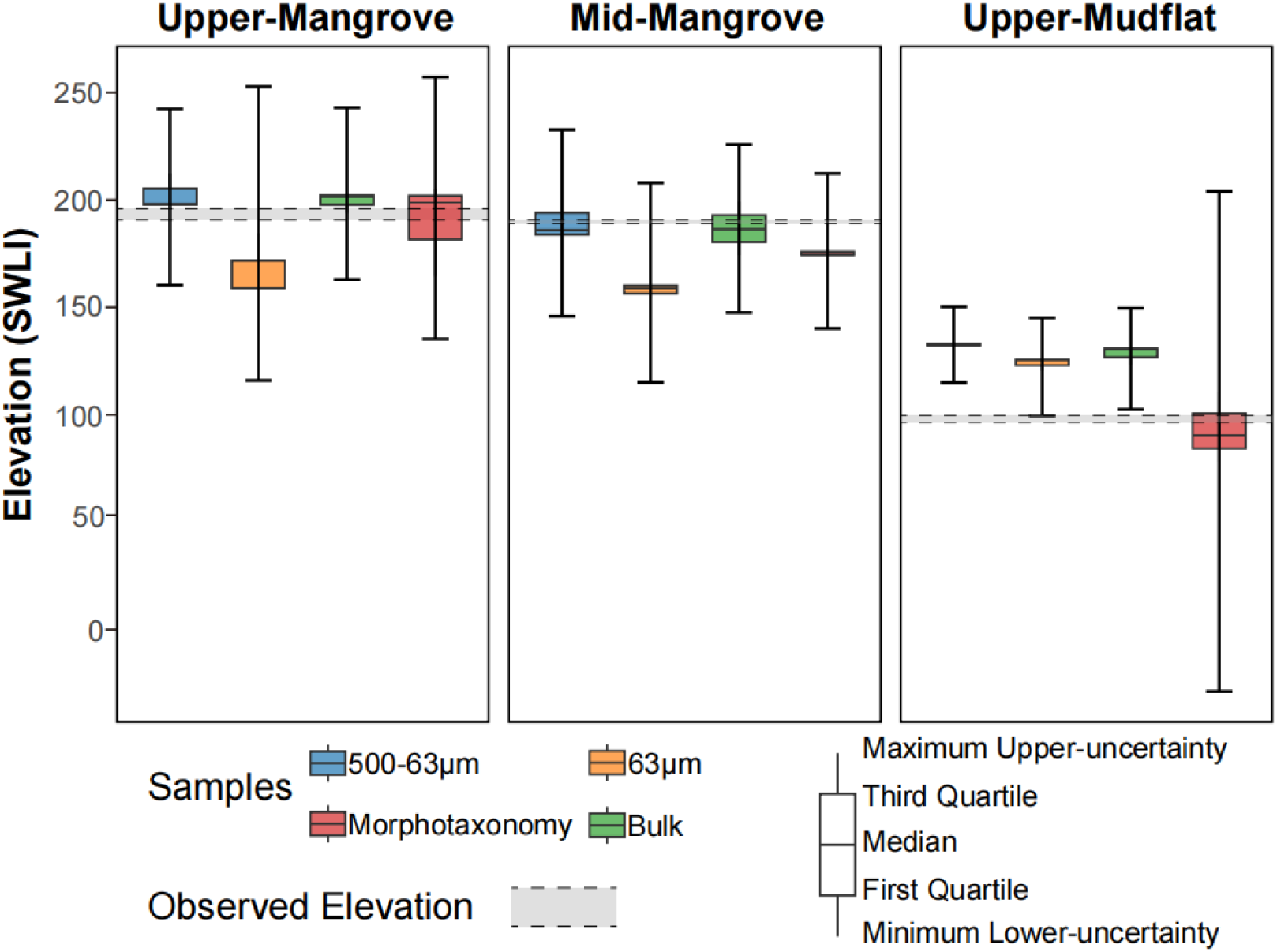
Elevation estimation (SWLI) of 500–63 μm, <63 μm size fractions and bulk samples of each sampling station produced by Bayesian transfer function (BTF) is depicted. The distributions of estimated SWLI for different size fractions of each station represent three replicate samples. The error bars indicate the maximum value of the 2-sigma uncertainty across these replicates. The observed elevation at each monitoring station is shown by the grey bar.

## 4. Discussion

### 4.1 eDNA sources of bulk sediment in different environments

The sources of foraminiferal eDNA from bulk sediment samples differ between mangrove and mudflat environments, reflecting variations in organic carbon content and life stages of present foraminifera. These contrasts suggest different dominant contributors to the bulk eDNA pool in mangrove versus mudflat settings, which could be fundamental for sea □level reconstructions based on bulk foraminiferal eDNA from samples originating in these environments. Mangrove environments integrate multiple eDNA sources, with relatively higher *in situ* contributions. In upper-mangrove sediments, bulk eDNA resembles the combined signals from both size fractions (500–63 □μm and <63 □μm) (PERANOVA; *p* >0.05), suggesting an integration of living adults of all sizes, juveniles, propagules, and DNA from dead specimens (Fig. 3A). Notably, when the overwhelming signal from Family Saccamminidae is excluded, the 500–63 □μm and bulk eDNA assemblages exhibit strong similarity (Fig. 3B), indicating a higher contribution from adult assemblages in bulk sediment eDNA at this station. In contrast, the mid-mangrove station displays distinct profiles for all three fractions (Fig. 3), including the 500–63 □μm fraction and bulk eDNA assemblage. It is because the bulk sediment eDNA reflects not only intracellular DNA from living and recently active individuals, but also accumulates a significant amount of extracellular eDNA (Nagler et al., 2022; Taberlet et al., 2018; Taberlet et al., 2012), which can represent >40% of total eDNA recovered from sediments (Ascher et al., 2009; Carini et al., 2016; Caro et al., 2023). It implies a potentially greater contribution of extracellular DNA to the bulk eDNA assemblage in mangrove environment. The high contribution of extracellular foraminiferal DNA in mangrove environment may be due to the high organic carbon content in mangrove soils and the resulting low pH (Liu et al., 2025b; Sun et al., 2021; Yu et al., 2025), which protects extracellular DNA and allows it to persist and accumulate over annual timescales (Angeles et al., 2020; Freeman et al., 2023; Schweizer, 2015). The high organic carbon content of mangrove sediments also implies slower degradation rates in this environment, including for extracellular DNA. Mangrove vegetation increases bottom friction and reduces tidal current velocities, resulting in greater flow damping compared to unvegetated mudflats (Wei et al., 2025; Zeng et al., 2025), which may create more sheltered conditions for dead organisms’ eDNA preservation. Similar to dead foraminiferal morphological assemblages, the accumulation of extracellular eDNA sources over longer time may cause bulk eDNA in these environments to reflect longer-term environmental signals, such as inundation frequency of tide (Corinaldesi et al., 2018; Nagler et al., 2022; Pedersen et al., 2015).

In contrast, mudflat environments predominantly reflect integrated signals from external sources, with a stronger imprint of temporally variable inputs. In the upper-mudflat station, bulk eDNA more closely resembles the <63 □μm fraction, indicating a predominance of propagule- and juvenile-derived DNA (Fig. 3A). This pattern likely results from multiple interacting factors: (1) lower organic carbon content reduces extracellular DNA preservation compared to mangrove environments (Sun et al., 2021; Yu et al., 2025), (2) high sedimentation rates that may rapidly bury and dilute local eDNA signals (Foster et al., 2024), and (3) greater transport of allochthonous material introduces propagule banks from deeper estuarine and marine environments (Singer et al., 2023), as supported by the higher relative abundance of taxa not observed in the morphological assemblages, such as the planktonic family Globigerinitidae (Fig. 2B). These combined processes favor the preservation of recent, transported DNA over accumulated local extracellular DNA, resulting in assemblages that integrate signals from multiple sources and more strongly reflect short □term temporal dynamics rather than long □term environmental conditions. These results are consistent with previous findings (Liu et al., 2025b), where eDNA assemblages from mangrove environments exhibited seasonal stability, while those from the upper-mangrove station were relatively unstable. Hence, there is a greater contribution from living adult populations in mangroves while in contrast, mudflat environments show pronounced temporal variation, likely driven by higher contributions of propagule-and juvenile-derived DNA. These environmental differences in the composition and sources of eDNA indicate that both the temporal resolution and ecological representativeness of eDNA-based RSL reconstructions may differ between mangrove and mudflat settings.

### 4.2 Implication for eDNA-based RSL reconstruction

To evaluate dead organisms-, propagule- and juvenile-derived eDNA’s influence on relative sea-level (RSL) reconstruction, we applied the Bayesian transfer function (BTF) developed in Liu et al (2025a) to estimate elevations from eDNA assemblages across size fractions and bulk samples at each station. Compared to morphological approaches, the eDNA-BTF overall demonstrated lower prediction uncertainty (Fig. 4), which aligned with previous study (Liu et al., 2025a). However, our analysis reveals environment-specific impacts: propagule and juvenile-derived eDNA significantly influenced elevation estimates in mudflat-mangrove transitional zone but exhibited minimal impact in mangrove environments.

Mudflat-mangrove transitional zone showed systematic biases across all fractions. In the upper-mudflat station, only the <63 μm fraction accurately predicted elevation within uncertainty limits, although all size fractions showed a systematic overprediction pattern (Fig. 4). The additional tested samples from the mudflat further showed that barren mudflat sediments can accurately reproduce elevation estimates, whereas both size fractions of the sample from the transitional zone were inaccurate and showed underprediction (Fig. S3). This inaccurate elevation estimate for the transitional zone likely results from two factors: (1) the transitional zone receives exogenous signals from both adjacent environments, and (2) high mobility and abundance of propagules in mudflats (Reef et al., 2017; Wang et al., 2023) broadens the range of elevation signals and skews results, particularly when Saccamminidae dominate the eDNA pool. The 500–63 μm fraction of upper-mudflat station, despite containing significantly fewer Saccamminidae sequences (*p* <0.05), remained influenced by high-intertidal taxa (e.g., OTU3 and OTU35), leading to overprediction. Mangrove environments demonstrated greater reliability. In contrast, both 500–63 μm and bulk eDNA from the upper- and mid-mangrove stations provided accurate elevation estimates. The stability and structure of the eDNA assemblage—bolstered by extracellular foraminiferal eDNA—support reliable reconstructions (Liu et al., 2025a; Liu et al., 2025b). While the mangrove <63 μm fraction still encompassed true elevations within its 2σ uncertainties, it showed greater uncertainty and systematic underprediction, potentially due to the reduced representation of high-elevation indicator taxa (e.g., Miliamminidae; Fig. 2) in the propagule pool during dry-season dormancy.

These environment-specific differences reflect how propagule-derived eDNA alters the taxonomic composition that determines elevation estimates. Given that abundant taxa dominate elevation signals (Fatela & Taborda, 2002; Walker & Cahill, 2024), propagule-dominated assemblages create prediction biases through two key mechanisms: the absence of key indicator taxa (as observed in mangrove <63 μm fractions) and the introduction of taxa from non-local elevations (as seen in mudflat environments). In mudflat-mangrove transitional zone, high propagule mobility and dominance create assemblages that poorly represent local elevation conditions, leading to systematic inaccurate prediction. Although barren mudflats are also dominated by high abundances of propagule-derived eDNA (Singer et al., 2023), these signals are not biased by inputs from different environments and therefore reflect more stable and accurate elevation signals. Conversely, in mangrove environments, extracellular DNA provides assemblage stability that maintains the taxonomic-elevation relationships essential for accurate RSL reconstruction. Therefore, eDNA-based reconstructions using bulk sediment are robust in mangrove settings, where extracellular DNA buffers against propagule-induced compositional changes. However, caution is warranted for mudflat-mangrove transitional zones, where dominance of propagules and the import of mixed exogenous DNA can fundamentally alter assemblage composition bias RSL estimation.

## 5. Conclusion

Our study highlights the influence of propagule and juvenile-derived DNA on foraminiferal eDNA assemblages and their application in relative sea-level (RSL) reconstruction across intertidal environments. By analyzing sieved and bulk sediment eDNA samples, we found that soft-walled monothalamids—particularly Saccamminidae—dominated the eDNA assemblages, especially in the <63 μm fraction, reflecting the significant contribution of propagules to the eDNA pool. While eDNA-based RSL reconstructions using a Bayesian transfer function (BTF) produced accurate elevation estimates for mangrove environments, including both upper- and mid-mangrove stations, the upper-mudflat station exhibited greater variability and overprediction, and <63 μm fraction in mangrove environment showed underprediction. This underscores the importance of considering propagule DNA contributions, particularly in mudflat–mangrove transitional zones, where exogenous inputs are more mixed and eDNA assemblages are more sensitive to environmental fluctuations. In line with previous studies, our results demonstrate that using foraminiferal eDNA assemblages extracted from bulk sediment are generally robust for RSL reconstruction in mangrove settings, with tidal elevation remaining the dominant shaping factor despite propagule-driven variations. However, caution should be taken when interpreting RSL reconstructions from mudflat-mangrove transitional zone samples, where propagule influence and exogenous inputs can reduce accuracy. Overall, our findings advance the understanding of foraminiferal eDNA dynamics and provide practical guidance for future paleoenvironmental reconstructions using eDNA-based proxies.

## Supporting information

Supplementary Figure - Propagule and Juvenile-derived Foraminiferal eDNA across intertidal habitats and its implications for accurate sea-level recons

Supplementary Table - Propagule and Juvenile-derived Foraminiferal eDNA across intertidal habitats and its implications for accurate sea-level reconst

## CRediT authorship contribution statement

**Liu Zhaojia**: Conceptualization, Data curation, Formal analysis, Methodology, Software, Visualization, Writing – original draft. **Nicole S. Khan**: Conceptualization, Funding acquisition, Project administration, Resources, Supervision, Writing – review and editing. **Magali Schweizer:** Conceptualization, Methodology, Writing – review and editing. **Celia Schunter**: Conceptualization, Resources, Supervision, Writing – review and editing.

## Acknowledgments

This research is supported by the Research Grants Council of Hong Kong (GRF Project no.: 27300221 and 17303925). The author would like to thank the World Wide Fund for Nature for granting permits for this study to conduct in the Mai Po Nature Reserve and their works in managing the natural reserve. Sincere appreciation to Gao Chengcheng and Qin Yonghui for their help in the field.

## Appendix A. Supplementary data

Supplementary Figures of this article can be found at file “Supplementary Figure - Environmental controls on propagule-derived foraminiferal eDNA and its implications for sea-level reconstruction”.

## Data Availability Statement

Raw sequence data were deposit in NCBI Sequence Read Archive under Bioproject: PRJNA1400784.

