## Supplementary Figure - Propagule and Juvenile-derived Foraminiferal eDNA across intertidal habitats and its implications for accurate sea-level recons for "Propagule and Juvenile-derived Foraminiferal eDNA across intertidal habitats and its implications for accurate sea-level reconstruction"

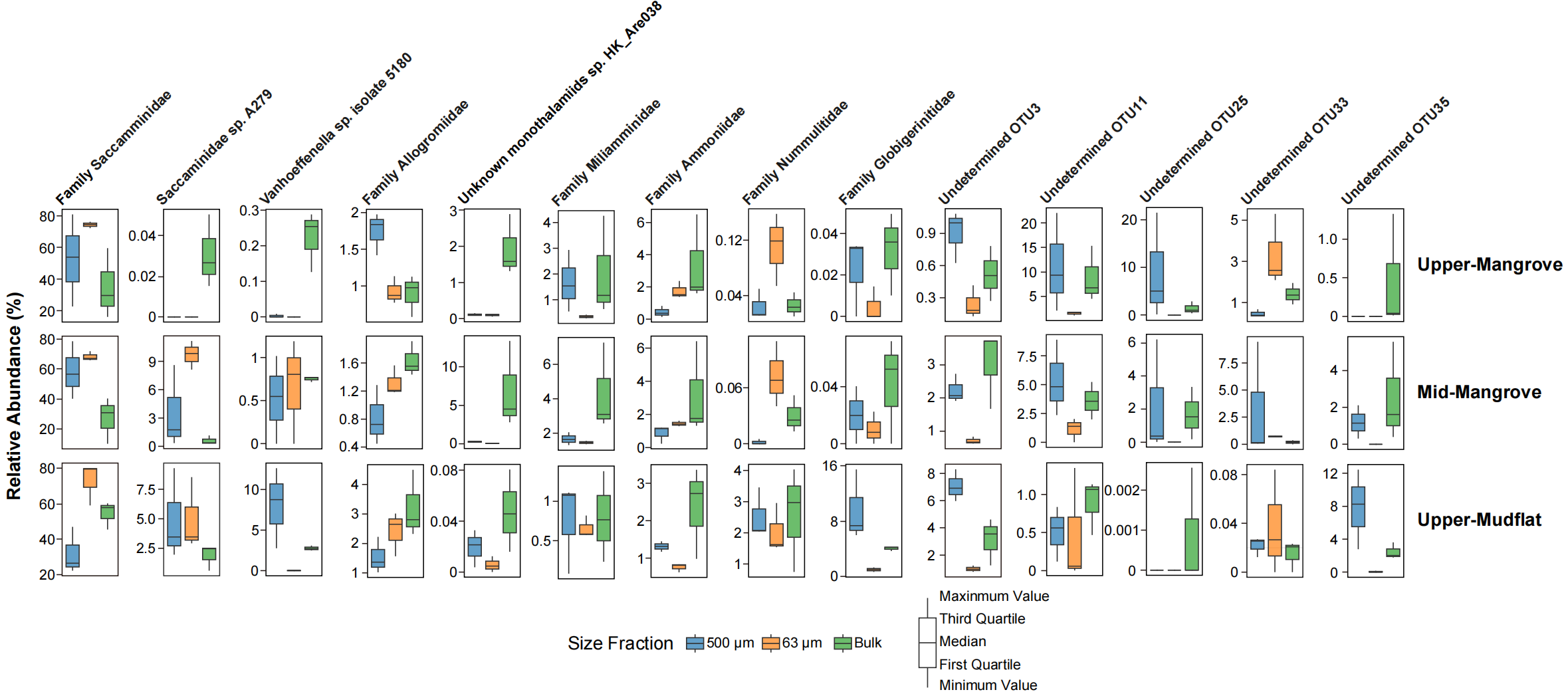

***Fig. S1*** Relative abundance of dominant taxa in 500–63 µm, <63 µm size fractions and bulk samples in foraminiferal eDNA assemblage at three sampling stations.

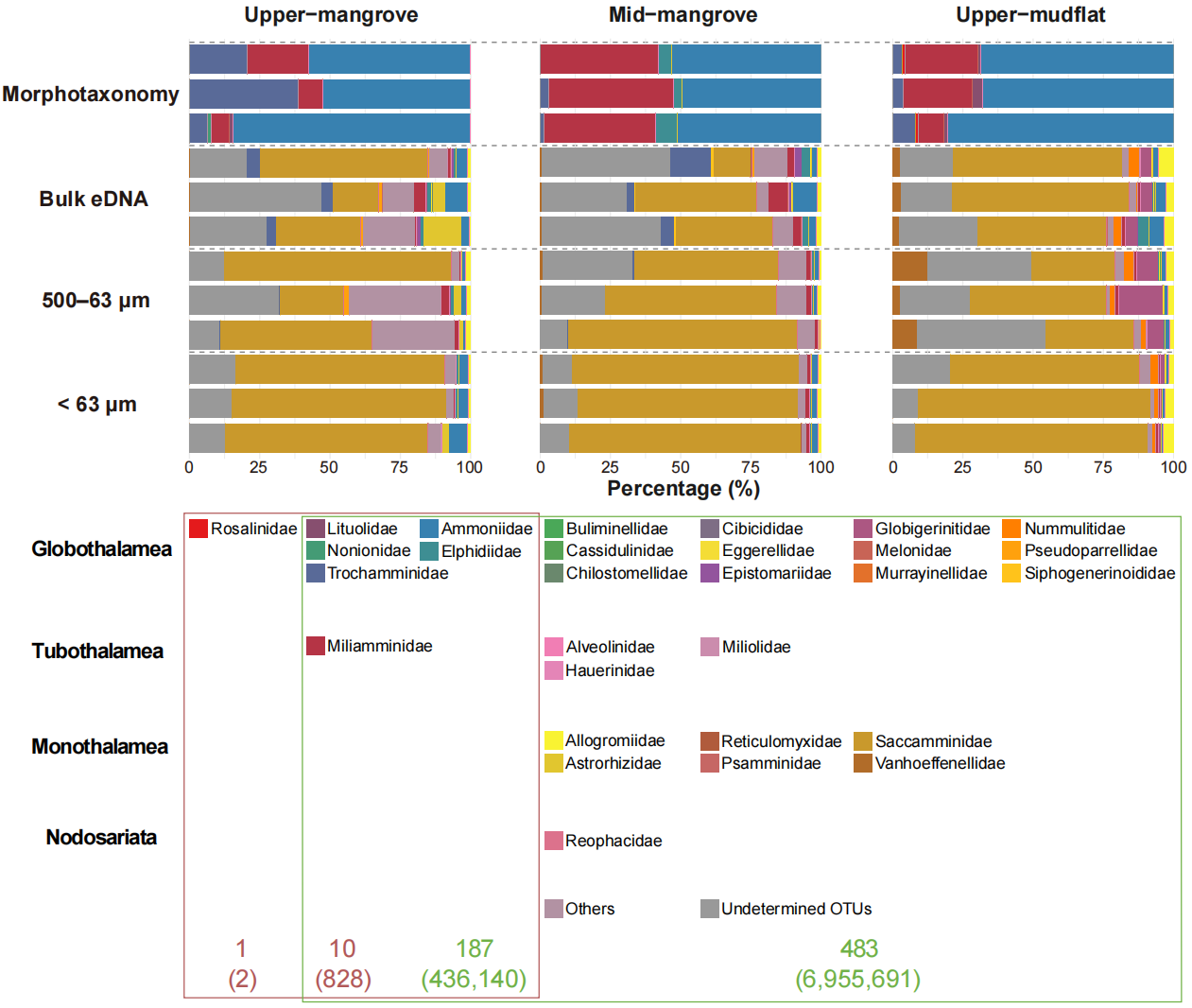
***Fig. S2*** Taxa abundance at Family level of morphological and eDNA assemblages of 500–63 µm, <63 µm size fractions and bulk samples in three sampling stations. The proportion of each taxon presented in the bar plot is related to its counts/reads number in the dataset. Red numbers correspond to morphological data, indicating the number of species (with the number of specimens in brackets). Green numbers correspond to molecular data, indicating the number of OTUs (with the number of sequences in brackets). “Others” includes OTUs that were assigned to order level.

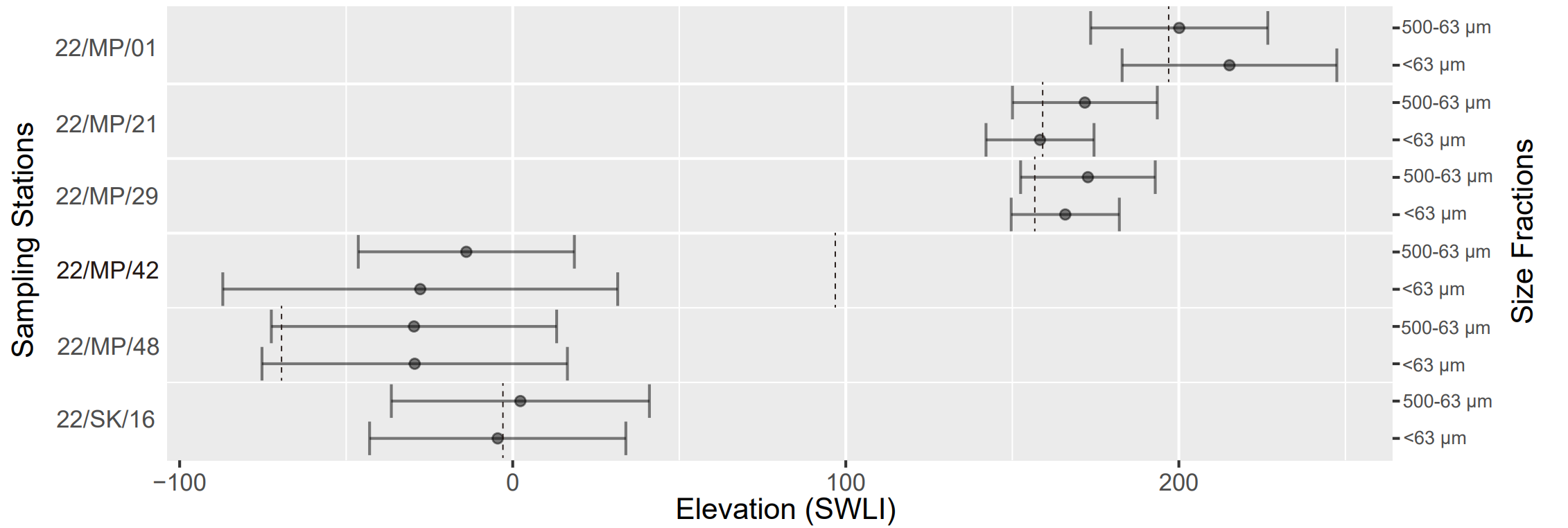

***Fig. S3*** Elevation estimation (SWLI) of 500–63 µm and <63 µm size fractions of samples collected at Liu et al (2025a) produced by Bayesian transfer function (BTF) are depicted. The error bars indicate the 2-sigma uncertainty of each estimation. The observed elevation at each sampling station is shown by the grey dash line.
